# A single checkpoint pathway eliminates mouse oocytes with dna damage or chromosome synapsis failure

**DOI:** 10.1101/137075

**Authors:** Vera D. Rinaldi, Ewelina Bolcun-Filas, Hiroshi Kogo, Hiroki Kurahashi, John C. Schimenti

## Abstract

Pairing and synapsis of homologous chromosomes during meiosis is crucial for producing genetically normal gametes, and is dependent upon repair of SPO11-induced double stranded breaks (DSBs) by homologous recombination. To prevent transmission of genetic defects, diverse organisms have evolved mechanisms to eliminate meiocytes containing unrepaired DSBs or unsynapsed chromosomes. Here, we show that the CHK2 (CHEK2)-dependent DNA damage checkpoint culls not only recombination-defective mouse oocytes, but also SPO11-deficient oocytes that are severely defective in homolog synapsis. The checkpoint is triggered by spontaneous DSBs that arise in late prophase I, accumulating above the checkpoint activation threshold (∼10 DSBs) because presence of HORMAD1/2 on unsynapsed chromosome axes prevents their repair. Furthermore, *Hormad2* deletion rescued fertility and meiotic DSB repair of oocytes containing a synapsis-proficient, non-crossover recombination defective mutation in a gene (*Trip13*) required for removal of HORMADs from synapsed chromosomes, indicating that a substantial fraction of meiotic DSBs are normally repaired by intersister recombination in mice.

---

Genetic and developmental analyses of mouse mutants have suggested there are at least two distinct checkpoints during meiotic prophase I in oocytes, one that monitors DSB repair, and another that monitors synapsis. Oocytes defective for either synapsis or DSB repair are eliminated with different dynamics and severity. Females with mutations causing pervasive asynapsis alone (e.g. *Spo11*^*-/-*^) are born with a severely reduced oocyte pool. The surviving oocytes undergo folliculogenesis but are reproductively inviable, becoming exhausted within a few weeks by atresia and ovulation ^1^. Oocytes defective in DSB repair alone (*Trip13*^*Gt/Gt*^), or defective in both synapsis and meiotic DSB repair (e.g. *Dmc1*^*-/-*^; *Msh5*^*-/-*^), are virtually completely eliminated between late gestation and wean age by the action of a DNA damage checkpoint ^1^,^2^. Furthermore, genetic ablation of meiotic DSB formation confers a *Spo11*^*-/-*^-like phenotype to such DSB repair mutants, consistent with the existence of separate DNA damage and synapsis checkpoints ^1^-^4^. For DSB repair, CHK2 (checkpoint kinase 2) signaling to TP53/TAp63 is required to eliminate *Trip13*^*Gt/Gt*^mutant oocytes that exhibit full chromosome synapsis but have unrepaired SPO11-induced DSBs ^5^. Interestingly, *Chk2* deficiency imparted a *Spo11* null-like phenotype upon *Dmc1*^*-/-*^ovaries, consistent with separate, sequentially-acting checkpoints ^5^. Genetic evidence for a distinct synapsis checkpoint came from studies of mice lacking HORMAD1 or HORMAD2, proteins which load onto axes of meiotic chromosomes throughout early prophase I, but are removed upon synapsis ^6^. Ablation of either in mice prevented loss of SPO11-deficient oocytes, resulting in the persistence of a [nonfertile] primordial follicle reserve in adults ^7^-^9^. These data suggested that the HORMADs are components of a synapsis checkpoint pathway. Another mechanism for elimination of oocytes, demonstrated for situations involving a single asynapsed chromosome, is related to the phenomenon of MSUC (meiotic silencing of unsynapsed chromatin), in which unsynapsed chromosomes or large unsynapsed chromosome segments are transcriptionally silenced, blocking progression past diplonema if those chromosomes include genes essential for oocyte survival and development ^10^.

Whereas these lines of evidence support the existence of separate checkpoints monitoring DNA damage and synapsis, studies in non-mammalian organisms indicate that the “pachytene checkpoint” – a term referring to delayed progression of meiosis or death of meiocytes triggered by genetic aberrations present in late pachynema – is more complex, consisting of both distinct and overlapping signaling pathways that also impact DNA repair modalities such as choice of recombination partner for the repair of meiotic DSBs (e.g. sister chromatid vs. homolog) ^11^-^14^. Here, we report the results of a series of experiments designed to discriminate whether the pachytene checkpoint in mouse oocytes indeed consists of distinct pathways responding to different signals, or if the responses are integrated into a single checkpoint pathway. Using a variety of mouse mutants defective for synapsis and/or DSB repair, coupled with mutants in putative pachytene checkpoint components, we present evidence that a canonical, CHK2- dependent DNA damage signaling pathway is responsible for eliminating oocytes that are defective in homologous recombination (HR) repair of DSBs, or which are highly defective for chromosome synapsis, a condition that leads to DSBs and CHK2- dependent elimination. Additionally, studies characterizing the dynamics of genetically- and environmentally-induced DNA damage indicate that the HORMAD proteins play a key role in eliminating defective oocytes by virtue of inhibiting the repair of SPO11- dependent and independent DSBs, likely by presenting a barrier to homologous recombination (HR) repair between sister chromatids, thus promoting DNA damage signaling. Taken together, we propose that the “pachytene checkpoint” operates on the basis of DNA damage signaling pathway and that extensive asynapsis leads to oocyte loss by inhibiting HR repair rather than triggering a distinct “synapsis checkpoint”.

## RESULTS

### CHK2 is required for efficient elimination of *Spo11*^*-/-*^oocytes

To investigate potential overlap in the meiotic DSB repair and synapsis checkpoint pathways in mice, we tested whether CHK2, a well-defined DSB signal transducer, contributes to the elimination of *Spo11*^*-/-*^oocytes that are asynaptic due to lack of programmed meiotic DSBs needed for recombination-driven homolog pairing.

Consistent with prior reports ^1^,^15^, we observed a greatly reduced number of total follicles in 3 week postpartum (pp) *Spo11*^*-/-*^ovaries compared to WT, and in particular, the oocyte reserve (pool of primordial resting follicles) was almost completely exhausted by 8 weeks of age (Fig. 1). Surprisingly, *Chk2* deletion dramatically rescued the oocyte reserve (Fig. 1a,b), albeit not to WT levels. The rescued follicles in double mutant females persisted robustly at least until 6 months pp (in one case, 554 total in a single ovary).

**Figure 1:**
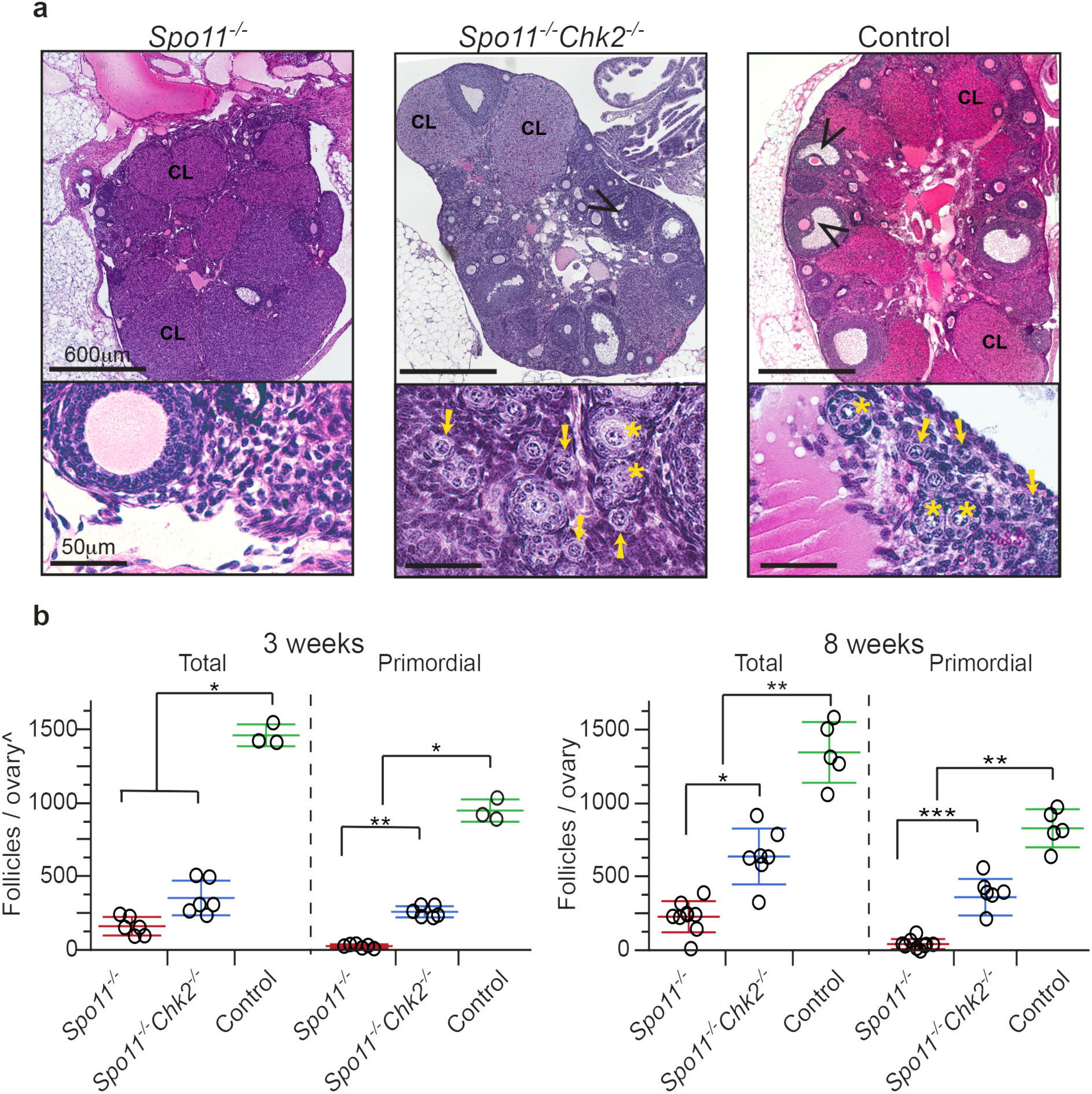
CHK2 is required for efficient elimination of asynaptic *Spo11*^*-/-*^mouse oocytes. **(a)** H&E stained histological sections of 8 weeks old ovaries. Black arrowheads indicate antral follicles. CL= Corpus Luteum; the presence of corpora lutea are indicate of prior rounds of ovulation. The lower portion of each panel contains a higher magnification image of an ovarian cortical region, where primordial follicles primarily reside. Yellow arrows and stars indicate primordial and primary follicles, respectively. **(b)** Follicle counts from ovaries of indicated genotypes at 3 and 8 weeks postpartum, respectively. Each data point is from a single ovary, each being from a different animal. Total = all follicle types. Horizontal hashes denote mean and standard deviation. ^ The values obtained for the 3 weeks follicles/ovaries counts are not comparable to the 8 weeks (see methods). Asterisks indicate p-values: (*) 0.005 ≤ p- values ≤ 0.05, (**)0.001 ≤ p-values ≤ 0.005 and (***) p-values ≤ 0.001 derived from a non-parametric, one-way ANOVA test (Kruskal-Wallis).

### HORMAD2 deficiency prevents elimination of *Trip13* mutant oocytes that have complete synapsis but unrepaired meiotic DSBs, restoring female fertility

Taken alone, the rescue of *Spo11*^*-/-*^oocytes by *Chk2* deletion suggests that severe asynapsis leads to CHK2 activation and signaling to mediate oocyte elimination. This led us to postulate that either: 1) CHK2 is a common component of otherwise distinct synapsis and DNA damage checkpoints, or 2) that there is a single linear checkpoint pathway that responds to both asynapsis and DNA damage, and that DNA damage activates the checkpoint pathway more robustly or sooner in prophase I (thus accounting for the different patterns of oocyte elimination in asynaptic vs. DSB repair-deficient oocytes mentioned above ^1^).

We reasoned that if there is a single linear checkpoint pathway, then putative synapsis checkpoint genes required to eliminate *Spo11*^*-/-*^oocytes would also be required to eliminate *Trip13*^*Gt/Gt*^oocytes. *Trip13*^*Gt/Gt*^meiocytes have synapsed chromosomes and persistent SPO11-dependent DSBs, which leads to neonatal depletion of follicles in a CHK2>TP53/TAp63 pathway-dependent manner (Fig. 2a)^2^,^5^. To test this, we determined whether deficiency of HORMAD2, a putative synapsis checkpoint protein, could rescue *Trip13*^*Gt/Gt*^oocytes. HORMAD2 and its paralog HORMAD1 are “HORMA” (<u>Ho</u>p1, <u>R</u>ev7 and <u>Ma</u>d2) domain-containing proteins orthologous to the *Saccharomyces cerevisiae* synaptonemal complex (SC) axial element protein Hop1p, and deletion of either prevents elimination of *Spo11*^*-/-*^oocytes ^7-^9. We used a mutant of *Hormad2* rather than *Hormad1*, because deletion of the latter disrupts homolog synapsis and DSB repair ^7^,^16^,^17^. Remarkably, not only did ovaries of 2 month old *Trip13*^*Gt/Gt*^ *Hormad2*^*-/-*^mice retain a substantial primordial follicle pool (Fig. 2a,b), but also these females were fertile (Fig. 2c). The rescued fertility of these oocytes suggested either that these DSBs were compatible with further oocyte maturation, or that they were eventually repaired as in the case of *Trip13*^*Gt/Gt*^females whose fertility was restored by *Chk2* ablation ^5^. The dynamics of DSB repair are addressed below.

**Figure 2:**
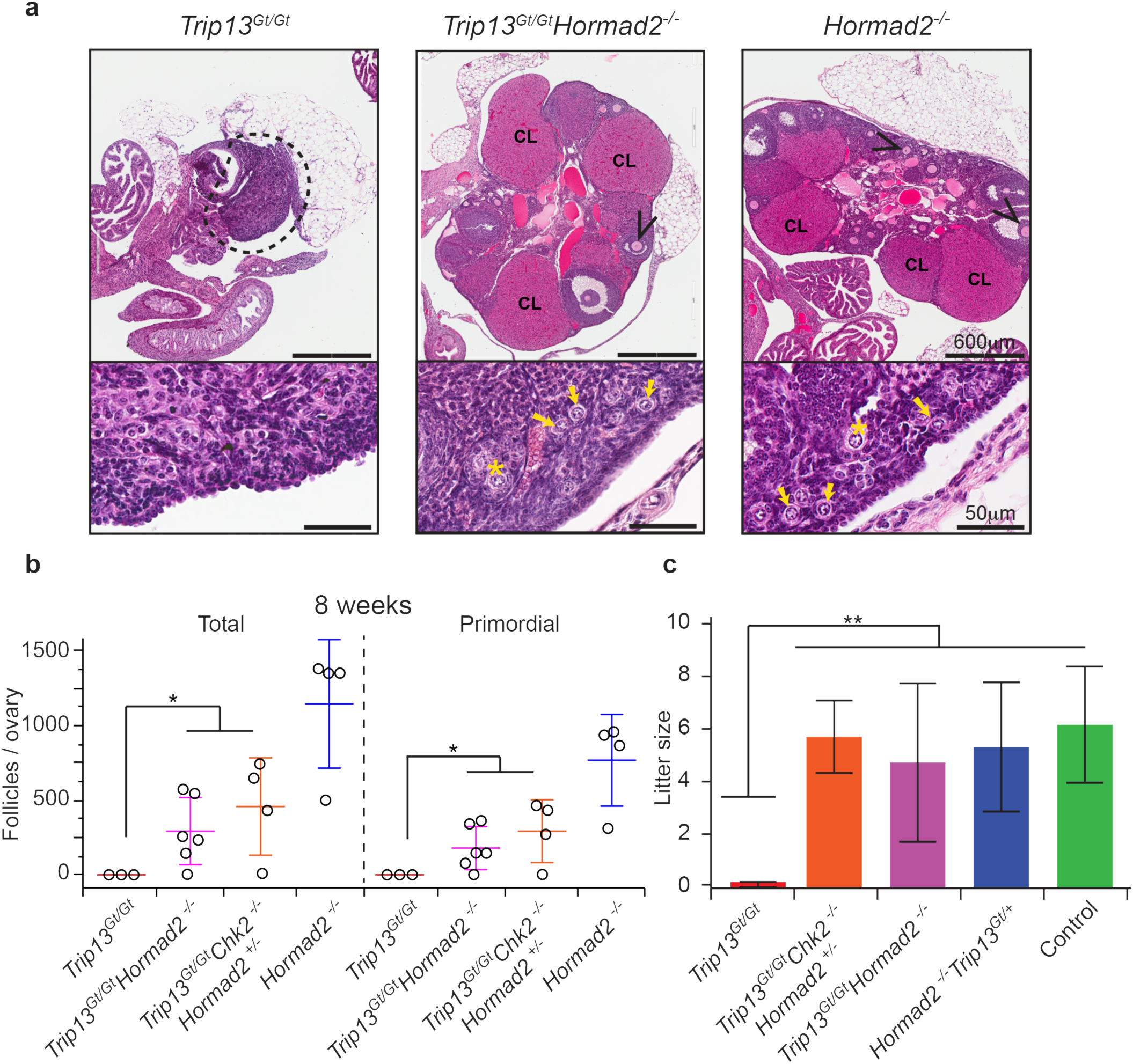
Synapsis-competent *Trip13*^*Gt/Gt*^oocytes are eliminated in a HORMAD2- dependent manner. (**a**) H&E stained histological sections of 8 week old ovaries of indicated genotypes. Black arrowheads indicate antral follicles. CL= Corpus Luteum. The lower half of each panel shows a higher magnification of cortical regions of ovaries. Yellow arrows and stars indicate primordial and primary follicles, respectively. (**b**) Follicle quantification of 8 week old ovaries. Each data point is from a single ovary, each being from a different animal. “Total” = all follicle types. Horizontal hashes denote mean and standard deviation. The statistic used was Kruskal-Wallis. * indicates p-value = 0.002. (**c**) Graphed are mean litter sizes. N ≥ 3 females tested for fertility per genotypic group. Error bars represent standard deviation and ** indicates p-value ≤ 0.005 derived from the Kruskal-Wallis test.

Since TRIP13 is required for depletion of the HORMADs from chromosome axes upon synapsis ^6^, and persistence of HORMADs on unsynapsed chromosomes correlates with MSUC-mediated silencing of essential genes ^8^,^10^, the question arises as to whether *Trip13*^*Gt/Gt*^oocytes are eliminated not because of unrepaired DSBs, but rather by transcriptional silencing. However, this is unlikely for the following reasons. First, *Trip13*^*Gt/Gt*^oocytes are depleted with a temporal pattern and degree consistent with mutants defective in DSB repair, not asynapsis ^1^,^2^. Second, *Spo11* is epistatic to *Trip13*, in that *Trip13*^*Gt/Gt*^ *Spo11*^*-/-*^ovaries resemble *Spo11* single mutants in their pattern of oocyte elimination ^2^, demonstrating that unrepaired meiotic DSBs drive early culling of *Trip13* mutant oocytes. Third, HORMAD persistence on synapsed *Trip13*^*Gt/Gt*^or unsynapsed *Spo11*^*-/-*^meiotic chromosome axes is not affected by *Chk2* deletion (Supplementary Fig. 1), which might be predicted if CHK2 was rescuing either mutant class by disrupting the ability of HORMADs to signal asynapsis. The latter is further supported by the fact that CHK2 depletion does not interfere with MSCI (meiotic sex chromosome inactivation, which is mechanistically similar or identical to MSUC) in males ^18^, and that *Chk2*^*-/-*^mice are fertile unlike *Hormad1*^*-/-*^animals ^7^,^16^,^19^.

### HORMAD2 inhibits DSB repair in prophase I oocytes

That HORMAD2 deficiency could rescue both *Trip13*^*Gt/Gt*^and *Spo11*^*-/-*^oocytes is consistent with a single checkpoint capable of detecting both damaged DNA and asynapsed chromosomes. If there is indeed a single checkpoint pathway, then combined deficiency for CHK2 and HORMAD2 should rescue asynaptic and DSB repair-defective *Dmc1*^*-/-*^oocytes to the same degree as deficiency for either one alone. However, *Dmc1*^*-/-*^ *Chk2*^*-/-*^ *Hormad2*^*-/-*^females had ≥3 fold increase in primordial and total follicles compared to *Dmc1*^*-/-*^ *Hormad2*^*-/-*^or *Dmc1*^*-/-*^ *Chk2*^*-/-*^ovaries (Fig. 3a,b, and Supplementary Fig. 2). This lack of epistasis indicates that HORMAD2 and CHK2 are not functioning solely as members of a single linear checkpoint pathway sensing either or both asynapsis and DNA damage.

**Figure 3:**
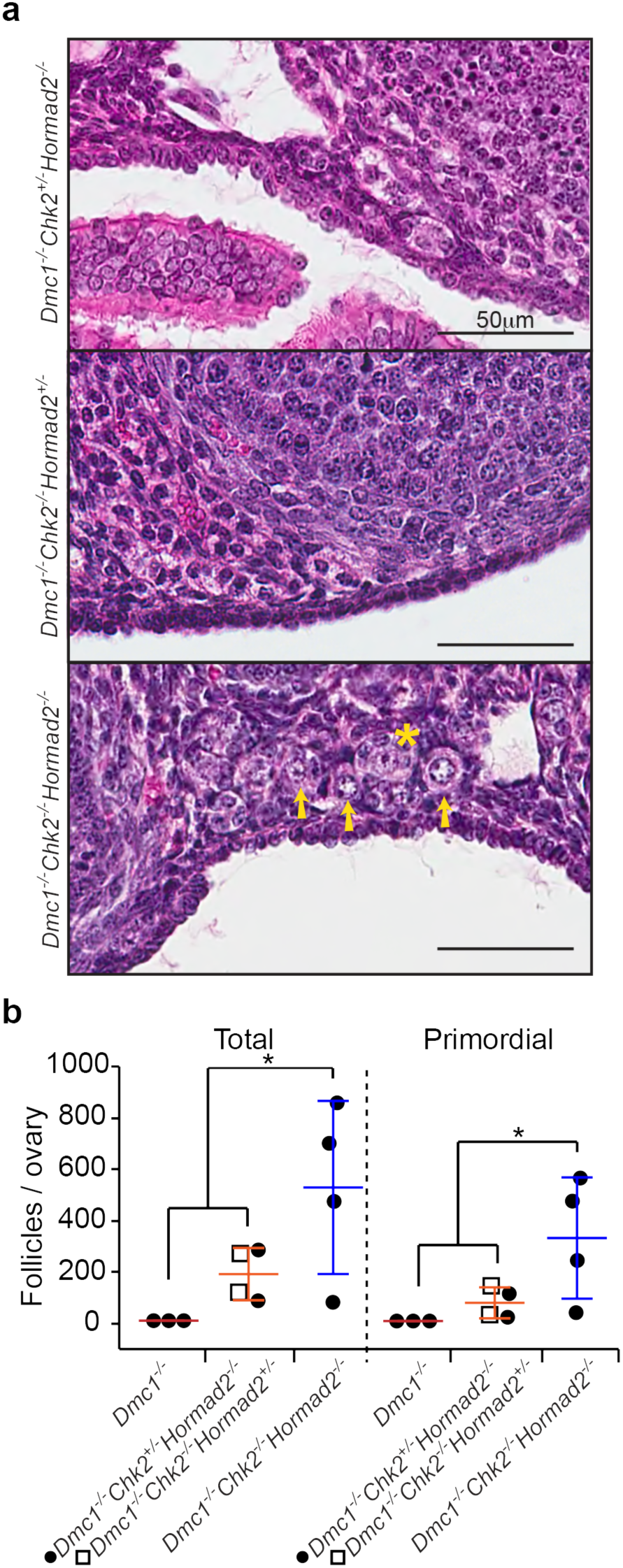
HORMAD2 and CHK2 are not in the same checkpoint pathway. (**a**) H&E stained histological sections of cortical regions of 8 week old mutant mouse ovaries, where primordial follicles are concentrated. Histology of whole ovaries of these genotypes are presented in Supplementary Fig. 1. Primordial follicles, which constitute the oocyte reserve, are indicated by yellow arrows, and a primary follicle by a star. Residual *Dmc1*^*-/-*^ovaries are not represented because they are completely devoid of oocytes ^54^. (**b**) Follicle counts from ovaries of indicated genotypes at 8 weeks of age. “Total” = all types of follicles. Data points represent follicle counts derived from one ovary, each ovary originating from a different animal. Asterisk indicates p-value ≤ 0.05 (Kruskal-Wallis test).

We therefore considered two alternative explanations for why *Hormad2* deficiency rescues *Trip13*^*Gt/Gt*^oocytes: 1) it reduces the number of SPO11-induced DSBs to a level sufficient for synapsis, but below the threshold for checkpoint activation; and/or 2) it promotes DSB repair. Studies of related proteins support both explanations. Absence of the budding yeast ortholog Hop1p, which is enriched together with Spo11p accessory proteins at locations of recombination hotspots, not only decreases meiotic DSB formation, but also increases use of the sister chromatid as a template for HR repair ^20-^25. Mouse spermatocytes lacking HORMAD1, which is required for loading its paralog

HORMAD2 onto unsynapsed axes and for normal SC formation ^7^, form >2 fold fewer SPO11-catalyzed DSBs ^7^,^26^, and is required for proper localization of MEI4 (a protein involved in meiotic DSB formation) to chromosome axes ^27^. However, although *Dmc1*^*-/-*^ *Hormad1*^*-/-*^or irradiated *Hormad1*^*-/-*^oocytes exhibit fewer DSB markers than oocytes containing HORMAD1 ^7^,^17^, this reduction in DNA damage appears to be attributable not only to decreased DSB formation, but also enhanced repair ^19^. Notably, an allele of the *C. elegans* HORMA protein HTP-3, proficient for DSB formation, revealed its role in promoting meiotic DSB repair in the absence of synapsis ^28^. Intersister (IS) HR repair of DSBs in *S. cerevisiae* is substantial and it increases in *hop1* mutants ^29^. Moreover, disruption of SC axes in mice (deletion of *Sycp2* or *Sycp3*) appears to alter recombination partner choice in favor of the sister chromatid, decreasing persistent DSBs in *Trip13*^*Gt/Gt*^oocytes to a degree that diminishes their elimination in a RAD54- dependent manner ^30^. These data led us to hypothesize that the rescue of *Trip13* mutant oocytes by *Hormad2* deficiency was due to enhanced IS HR repair of DSBs, suggesting that the HORMADs normally inhibit IS repair until they are removed from prophase I chromosomes upon synapsis.

To test this, we quantified levels and rates of meiotic DSB repair in various genotypes of prophase I oocytes. Whereas the number of leptotene and zygotene stage RAD51 foci was not significantly different in *Trip13*^*Gt/Gt*^ *Hormad2*^*-/-*^oocytes compared to *Trip13*^*Gt/Gt*^or other control and mutant genotypes (Fig. 4a,b), there were significantly fewer compared to *Trip13*^*Gt/Gt*^by pachynema, and nearly all RAD51 foci disappeared by diplonema (to a level not statistically different from WT or *Hormad2*^*-/-*^oocytes). In contrast, RAD51 levels in *Trip13*^*Gt/Gt*^and *Trip13*^*Gt/Gt*^ *Chk2*^*-/-*^(newborn) oocytes remained high in diplonema (Fig. 4a,b). Furthermore, we found that RAD51 foci induced by 2Gy of ionizing radiation (IR) disappeared more rapidly (8 hrs post-IR) in *Spo11*^*-/-*^ *Hormad2*^*-/-*^oocytes than either *Spo11*^*-/-*^or *Spo11*^*-/-*^ *Chk2*^*-/-*^oocytes (Fig. 5). Overall, the data indicate that HORMAD2 on the axes of either asynapsed (*Spo11*^*-/-*^) or synapsed (*Trip13*^*Gt/Gt*^) ^6^ meiotic chromosomes inhibits DSB repair, most likely via IS recombination.

**Figure 4:**
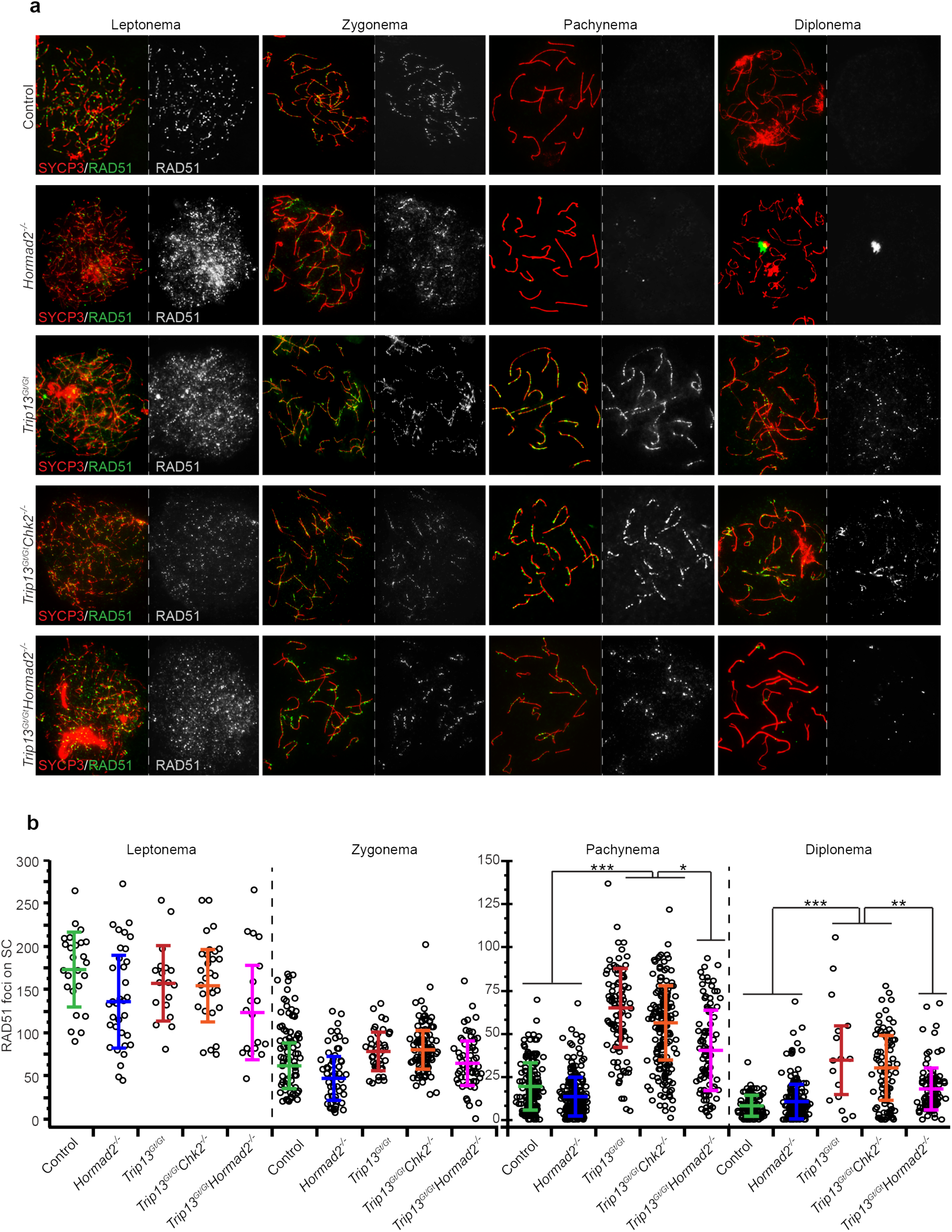
Depletion of HORMAD2 accelerates DSB-repair during early stages of meiotic prophase I. (**a**) Representative images of meiotic chromosome spreads from oocytes at substages of meiotic prophase I, probed with antibodies for SYCP3 (SC axis protein) and the DSB marker RAD51. Oocytes were isolated from female embryos ranging from 15.5 dpc to newborns. (**b**) Numbers of RAD51foci in meiotic prophase I substages of mutants. Only RAD51 foci present on SYCP3 stained axes were scored. Each data point represents one cell. In each genotypic group, at each stage, the counts are derived from at least three animals. Horizontal hashes in summary statistic plots denote mean and standard deviation. Colors correspond to genotypes. Asterisks indicate statistical significant differences between groups in terms of the least square means of RAD51 foci. p-values: *** p ≤ 0.001; ** p ≤ 0.005; * p ≤ 0.05 (Tukey HSD).

**Figure 5:**
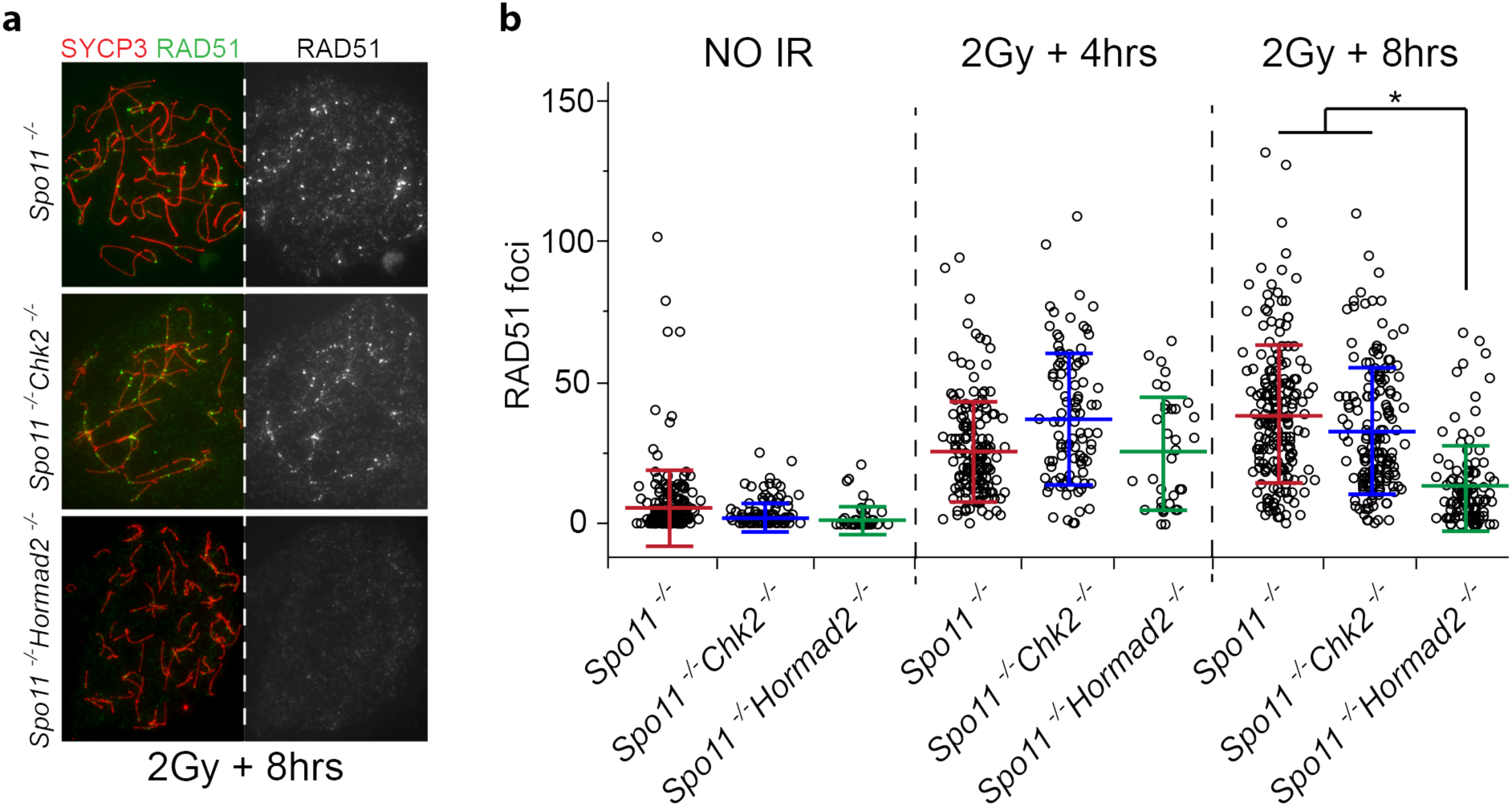
Depletion of HORMAD2 accelerates repair of induced DSBs in oocytes. Fetal ovaries were collected at 15.5dpc, cultured 24 hours, exposed to 2 Gy of ionizing radiation (IR), then cultured for an additional 4-8 hours. (**a**) Immunolabeling of surface spread chromosomes from oocytes collected 8hrs after IR. (**b**) Quantification of RAD51 foci. Each data point represents one oocyte. The graphs include mean and standard deviation, and are color coded according to genotypic group. The 4 and 8 hr unirradiated samples were combined. Data were derived from at least two different animals per condition.

## Evidence that CHK2-mediated elimination of asynaptic oocytes is driven by accumulation of SPO11-independent DSBs

If indeed *Hormad2* deletion rescues DSB-containing oocytes by weakening or eliminating a barrier to sister chromatid recombination (BSCR), this raises the question as to why HORMAD2 deficiency rescues *Spo11*^*-/-*^oocytes that don’t make meiotic DSBs. A clue comes from the surprising observation that *Spo11*^*-/-*^oocytes sustain DSBs of unknown origin (but possibly from LINE-1 retrotransposon activation {Malki, 2014 #4699}) during early pachynema ^31^. We hypothesized that these DSBs occur at levels sufficient to trigger the CHK2-dependent checkpoint in *Spo11*^*-/-*^oocytes, but that in the absence of HORMAD2 there is sufficient DSB repair to prevent checkpoint activation. To test this, we determined the threshold number of DSBs that kills WT and *Chk2*^*-/-*^oocytes by exposing explanted newborn ovaries to a range of IR. RAD51 foci accumulated roughly linearly in oocytes exposed to 0.5 - 9Gy (Fig. 6a), and *Chk2*^*-/-*^oocytes withstood up to 7Gy (Fig. 6b), a dosage that induces ∼73.3 RAD51 foci (Fig. 6a). In contrast, as little as 0.3Gy (10.3 foci by linear regression) abolished the entire primordial follicle pool of WT ovaries. The SC axes of *Spo11*^*-/-*^zygotene/pachytene-like chromosomes in newborn oocytes contained an average of 30.4 discrete RAD51 foci (Fig. 6c; likely an underestimate, see Supplementary Fig. 3 and Discussion) indicating that majority of *Spo11*^*-/-*^oocytes (60.76% as compared to 29.05% in wild type) bear a level of DSBs (>10.3 foci) sufficient to trigger their elimination by the CHK2-dependent DNA damage checkpoint.

**Figure 6:**
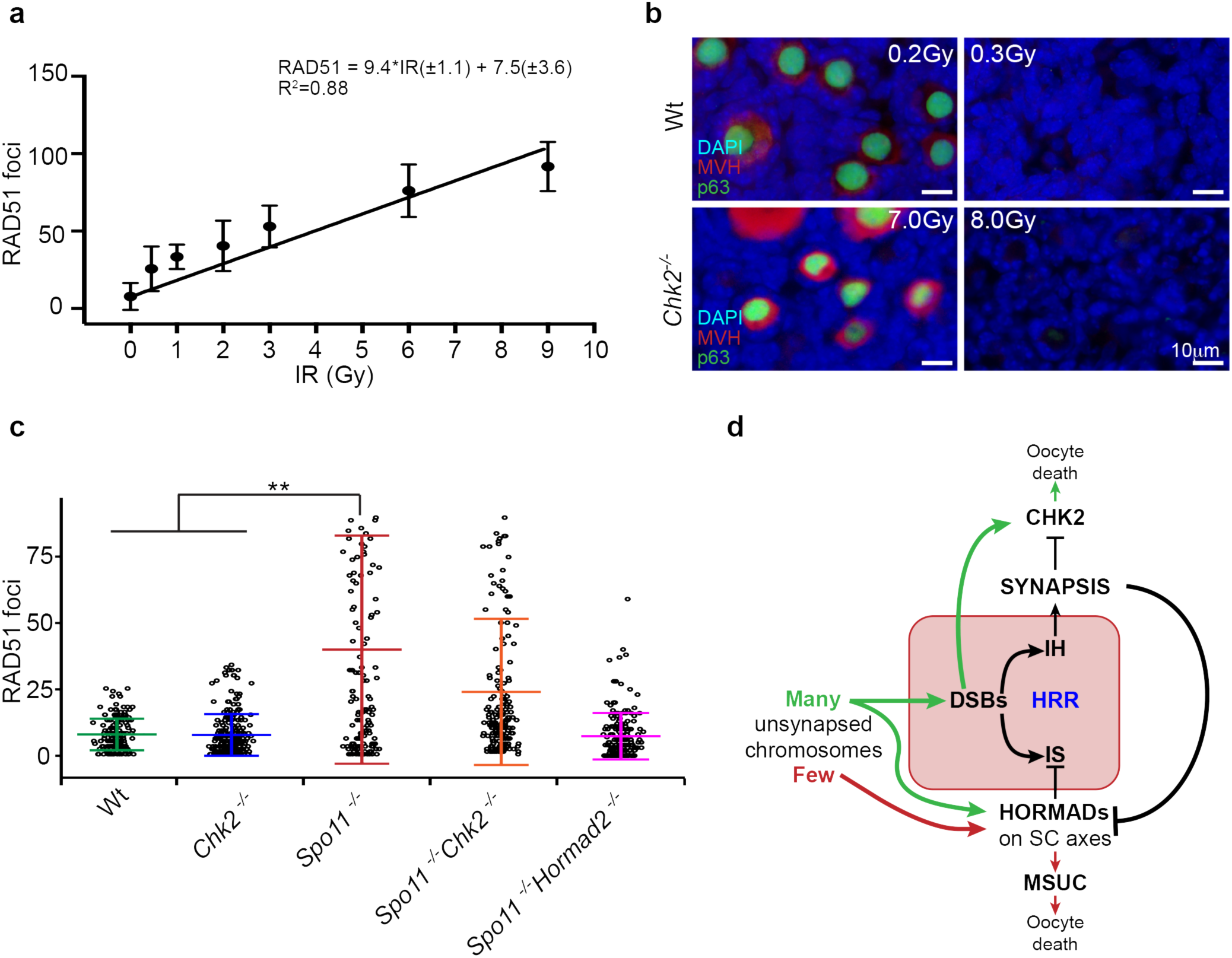
DNA damage threshold required to trigger oocyte death, and evidence for HORMAD-mediated inhibition of IS repair. (**a**) Linear regression for conversion of radiation dosages to RAD51 focus counts. Meiotic surface spreads were made from WT neonatal ovaries 2.5 hrs after IR. Plotted are means with standard deviations. Each IR dose has focus counts from at least 30 oocytes derived from at least two different animals (18 total). (**b**) *Chk2*^*-/-*^oocytes are highly IR resistant. Shown are immunofluorescence images of ovarian sections labeled with nuclear and cytoplasmic germ cell markers (p63 and MVH, respectively). (**c**) RAD51 focus foci counts from newborn oocyte spreads. Only oocytes with discrete patterns of RAD51 foci were scored, as defined in Supplementary Fig. 2. Data points represent individual oocytes, derived from at least five different animals from each genotypic group. Horizontal hashes denote means and standard deviations. Asterisks indicate statistically significant differences between groups with p-values: *** p ≤ 0.001; ** p ≤ 0.005; * p ≤ 0.05 (Tukey HSD). (**d**) Model for pachytene checkpoint activation in mouse oocytes. Oocytes with many unsynapsed chromosomes (green) ultimately accumulate DSBs, which cannot be repaired due to block to IS recombination imposed by HORMADs on asynapsed axes. Failure of DSB repair leads to activation of CHK2 and downstream effector proteins (p53/TAp63) that trigger apoptosis. Few asynapsed chromosomes (red) lead to inactivation of essential genes by MSUC thereby causing oocyte death. HRR -Homologous Recombination Repair; IH - Interhomolog; IS - Intersister; MSUC - Meiotic Silencing of Unsynapsed Chromatin.

## Discussion

Genome maintenance in germ cells is critical for fertility, prevention of birth defects, and the genetic stability of species. Throughout mammalian germ lineage development, from primordial germ cells (PGCs) through completion of meiosis, there are mechanisms that prevent transmission of gametes with genetic defects. Indeed, mutation rates in germ cells are far lower than in somatic cells ^32^-^34^. This is reflected by the exquisite sensitivity of PGCs to mutations in certain DNA repair genes ^35^-^37 38^, resting oocytes to clastogens such as radiation and chemotherapeutics ^39^-^41^, and developing prophase I meiocytes to genetic defects including a modicum of DNA damage ^41^,^42^ (Fig. 6b) or the presence of single asynapsed chromosome or even a chromosomal subregion ^43^,^44^.

Checkpoint responses of meiocytes entail adaptations to their unique developmental circumstances, chromosome biology and cell cycle. For example, mechanisms must exist to ensure that a pachytene/prophase I checkpoint is active only at a point in prophase I at which DSBs have normally been repaired, but not during the time between programmed DSB formation and HR repair, a period of several days in both oocytes and spermatocytes. It is noteworthy that in prophase I cells, quality control checkpoints are formally occurring in M phase (by analogy to mitotic cells). Mitotic cells do not have an intra-M checkpoint, but rather a G2/M checkpoint. However, oocytes that have progressed through prophase I to the dictyate stage are generally considered to be in G2-like arrest stage ^45^.

While the oocyte "pachytene checkpoint" is distinct with respect to its cell cycle timing and its ability to monitor an event (chromosome synapsis) unique to meiosis, our prior ^5^ and current work indicate that for circumstances involving extensive asynapsis and DNA damage, this checkpoint in oocytes consists of a DNA damage response common to somatic cells. Our surprising finding that culling of *Spo11*^*-/-*^oocytes occurs via CHK2 signaling raises the question of how SPO11-independent DSBs - first reported by Carofiglio *et al* ^31^ and confirmed here - arise on unsynapsed chromosomes. One possible source is LINE-1 retrotransposon activation, which has been correlated with natural oocyte attrition ^46^. However, transposon expression normally occurs only transiently at the onset of meiosis before epigenetic silencing ^47^. It is possible that the extensive asynapsis in *Spo11*^*-/-*^oocytes *per se*, or disruption of the meiotic program including the normal course of DSB induction and repair, interferes with transposon silencing. Another possibility is that unsynapsed chromosomes are more susceptible to spontaneous breakage. These outcomes could be exacerbated by extended retention of HORMADs on unsynapsed axes, all while inhibiting IS repair of newly arising DSBs. An intriguing question is whether the production of these SPO11-independent DSBs, whatever their origin, evolved as a contributory mechanism for genetic quality control. It is also conceivable that the extended presence of HORMADs themselves contributes to spontaneous DSB formation, possibly as a "last ditch" mechanism to drive pairing or synapsis in chromosomes devoid of sufficient interhomolog recombination events.

The late appearance and highly variable number (Fig. 6c) of SPO11-independent DSBs in *Spo11*^*-/-*^oocytes may explain the differences in timing and extent of oocyte elimination in exclusively asynaptic *vs.* DSB repair-deficient (e.g. *Dmc1*, *Trip13*) mutants. As reported by Di Giacomo and colleagues ^1^, whereas *Dmc1*^*-/-*^oocytes were completely eliminated before dictyate arrest and follicle formation, *Spo11*^*-/-*^ovaries contained ∼15-20% of WT numbers of follicles (including 27 fold less primordial follicles by 4 days pp); this reduced oocyte reserve was depleted by 2-3 months of age by subsequent cycles of recruitment and maturation. Additionally, *Dmc1*^*-/-*^oocytes degenerated before *Spo11*^*-/-*^oocytes, suggesting that an earlier-acting mechanism was triggering *Dmc1*^*-/-*^oocyte death. These distinctions, in conjunction with epistasis analysis of mutants doubly deficient for *Spo11* and DSB repair mutations, led to the conclusion that there are DSB-dependent and -independent mechanisms to eliminate defective oocytes. We suggest a modification to this interpretation: that the DSB-dependent mechanism (the DNA damage checkpoint) eliminates most oocytes of both asynaptic and DNA-repair deficient genotypes, but at different times. Since abundant programmed SPO11 DSBs occur early in prophase I, we posit that these trigger the checkpoint sooner and more uniformly in DSB repair mutants than in *Spo11*^*-/-*^oocytes, in which spontaneously DSBs occur in late prophase I, leading to later onset of apoptosis in those with threshold levels of DSBs. Based on our data (Fig. 6a), we suggest that those oocytes with below-threshold DSB levels escape the checkpoint, thus surviving to constitute the reduced follicular reserve in *Spo11* mutants.

While the CHK2-dependent checkpoint is of central importance to genetic quality control in oocytes, our observations that *Chk2* deletion does not fully restore oocyte numbers to WT levels in mutants indicates that it is not absolutely required for eliminating all oocytes with unrepaired DSBs. Rather, the fraction of oocytes rescued is inversely related to the burden of unrepaired meiotic DSBs. For example, whereas *Chk2* deficiency rescued nearly 1/3 of *Trip13*^*Gt/Gt*^oocytes (which are partially proficient for DSB repair and which harbor 35±4 and 63±4.7 persistent RAD51 foci in diplonema and pachynema, respectively; Fig. 4b), it rescued only a small fraction (∼5%) of profoundly recombination-deficient *Dmc1*^*-/-*^oocytes (harboring an average of ∼150 RAD51 foci ^30^). We posit that the oocytes that fail to be rescued in these mutants are eliminated either by another checkpoint pathway, or succumb from catastrophic levels of DNA damage. It is informative that deletion of *Hormad1*, but not *Hormad2*, rescues *Dmc1*^*-/-*^oocytes to a greater extent than *Chk2* deletion. As discussed earlier, the rescued *Dmc1*^*-/-*^ *Hormad1*^*-/-*^oocytes had a marked reduction in DSBs ^5^,^8^,^19^. Since HORMAD1 is needed to load HORMAD2 onto unsynapsed chromosome axes (not vice versa), then the impact of *Hormad1* deletion upon IS recombination constitutes the combined roles of both HORMAD proteins. However, when *Hormad2* alone is deleted, the continued presence of chromosomally-bound HORMAD1 provides a less-effective, but still substantive, BSCR. The lower level of residual DSBs in *Spo11* and *Trip13* mutant oocytes (compared to *Dmc1*^*-/-*^) may render them responsive to a weaker BSCR such as when *Hormad2* is deleted. We postulate that because of its involvement in stimulating SPO11 activity ^7^, *Hormad1* deletion is very effective in rescuing a DSB repair mutant like *Dmc1* because not only are fewer DSBs formed, but also IS recombination is more active.

Our results add to increasing evidence that IS recombination, and modulation thereof, is critically important in mammalian meiosis. As discussed in the text, the HORMADs and SC axial element structure appear to inhibit IS repair of meiotic DSBs, thus allowing IH recombination to drive homolog pairing and synapsis. However, as synapsis progresses and the SC is formed, the HORMADs are removed and presumably IS recombination can occur. Since not all RAD51 foci disappear by pachynema when synapsis is complete (for example, see Fig. 4b), it is possible that these DSBs are destined for, or normally repaired by, IS recombination. We speculate that the persistent unrepaired DSBs on synapsed chromosomes of *Trip13* mutants, which retain HORMADs on their SCs, may actually constitute that fraction of SPO11- induced DSBs (an average of ∼65/oocyte nucleus; Fig. 4) that would normally be repaired by IS recombination. If so, this represents a major fraction of all meiotic DSBs (∼200-300) formed in mammalian meiocytes.

In trying to decipher the quality-control mechanisms functioning during meiosis, it is important to recognize that experimental studies such as those performed here employ mutants with pervasive, non-physiological levels of defects. Meiocytes in wild-type individuals would have less extreme genetic defects. For oocytes, two reports have provided insightful information in this regard ^10^,^48^. These studies found that in oocytes bearing a small number (1-3) of asynapsed chromosomes, arising spontaneously in WT mice (both studies), from mutations ^48^, or from breeding-induced aneuploidies ^10^, that those asynapsed chromosomes underwent transcriptional silencing (MSUC) during pachynema (characterized by BRCA1-induced phosphorylation of H2AX, lack of Pol II binding, and ubiquitination of H2A), leading to elimination at the diplotene stage.

Compelling evidence that the silencing of essential meiosis or housekeeping genes on these chromosomes was responsible for the oocyte elimination, rather than a canonical checkpoint, came from the finding that MSUC of supernumerary chromosomes did not lead to oocyte demise ^10^. However, Kouznetsova and colleagues found that MSUC was compromised in oocytes with more than 2-3 unsynapsed chromosomes, presumably due to a limiting amount of BRCA1 ^48^. This is seemingly difficult to reconcile with the phenomenon of “pseudo sex body” formation in a substantial fraction of *Spo11*^*-/-*^meiocytes, in which a small number of asynapsed autosomes undergoes apparent MSUC in a manner analogous to MSCI of the XY (sex) body ^49^. Formation of pseudo sex bodies in *Spo11*^*-/-*^oocytes is dependent upon HORMADs ^7^,^16^, leading to the proposal that this is involved in oocyte elimination ^9^. However, it would seem that unless the chromosomal region(s) affected in a pseudo sex body in an oocyte are haploinsufficient (individually or collectively so), or both alleles happen to be simultaneously silenced in an oocyte, this manifestation of MSUC would not likely affect a large fraction of completely asynaptic oocytes. Furthermore, since CHK2 deficiency rescues *Spo11*^*-/-*^oocytes but doesn’t abolish HORMAD localization (Supplementary Fig. 1) or pseudo sex body formation (not shown), this contradicts the idea that MSUC is the “checkpoint” that leads to death of oocytes with pervasive asynapsis. Finally, because MSUC involves many components of the DNA damage response ^50^-^52^, it is conceivable that asynapsis leading to MSUC would activate effector elements of the DNA damage checkpoint pathway, including CHK2. However, this does not appear to be the case, because (as mentioned above) silenced supernumerary chromosomes do not eliminate oocytes, MSCI does not kill spermatocytes, and asynaptic oocytes are not eliminated in a pattern typical of DNA repair mutants.

Overall, this study shows that in mice, the “pachytene checkpoint” – long thought to consist of separate DNA damage and synapsis checkpoints – actually consists of only one true checkpoint: the DNA damage checkpoint. We believe the current information supports a model (Fig. 6d) for two major mechanisms by which oocytes with synapsis defects are eliminated: 1) MSUC, for oocytes with a small number of asynapsed chromosomes that do not accumulate unrepaired DSBs above a threshold, and in which both homologs of chromosomes bearing essential genes for meiotic progression are silenced ^10^, and 2) the DNA damage checkpoint, for oocytes with multiple asynapsed chromosomes that accumulate a sufficient number of DSBs to trigger the DNA damage checkpoint (Fig. 6d). These disparate mechanisms may have distinct purposes.

Because oocytes with only 1 or 2 unsynapsed chromosomes may not efficiently trigger the spindle assembly checkpoint (SAC) ^53^, the MSUC pathway would safeguard against aneuploidy. Superficially, it would seem that because oocytes with extensive asynapsis would effectively trigger the SAC, that the DNA damage checkpoint mechanism is redundant. However, it is likely advantageous reproductively to eliminate such defective oocytes before they enter dictyate as constituents of the ovarian reserve, otherwise the fraction of unproductive ovulations (those terminated by the SAC) would increase, thus compromising fecundity.

## Acknowledgements

This work was supported by a grant from the National Institutes of Health (R01 GM45415 to JCS), and contract CO29155 from the NY State Stem Cell Program (NYSTEM). The authors would like to thank R. Munroe and C. Abratte for generating chimeric mice, Stephen Parry from the Cornell Statistical Consulting Unit (CSCU) for help with the statistical analysis, Dr. Atilla Toth for the HORMAD2 antibody, and M.A. Handel for feedback on the manuscript.

## MATERIALS and METHODS

### Animals

The alleles used have been previously described and were the following: *Trip13*^*Gt(RRB^047^)Byg*^(referred to as *Trip13*^*Gt*^in the manuscript) ^1^; *Dmc1*^*tm*^1^*Jcs*^^2^; *Chk2*^*tm*^1^*Mak*^^3^; *Spo11*^*tm*^1^*Sky*^^4^; and *Hormad2* ^5^. All mice were in a mixed genetic background of strains C57Bl/6J and C3H/HeJ. Comparisons between compound mutants and controls were done using littermates or related animals. Unless otherwise noted, all experiments used at least three mice per experimental group. The embryonic age was counted using the morning in which copulation plug was detected as being the 0.5 days post coitus (dpc). The Cornell’s Animal Care and Use Committee approved all animal usage, under protocol 2004-0038 to JCS.

### Organ Culture and Irradiation

Embryonic and postpartum ovaries were cultured as previously described ^6^-^8^, with the exception of the use of inserts with pore size of 8μm. Ovaries were irradiated in a 137cesium irradiator with a rotating turntable. Immediately after irradiation, the media was replaced, and ovaries were cultured for indicated periods of time prior to tissue processing. All the irradiation experiments reported here were done on explanted ovaries.

### Histology and Immunostaining

Ovaries were dissected and fixed in Bouin’s fixative overnight. Afterwards, tissues were washed in 70% ethanol prior to being embedded in paraffin for serial sections at 6μm thickness. Ovaries were stained with Harris Hematoxylin and Eosin (H&E) and follicles counted in every fifth section except for the three-week counts reported in Figure 1b, in which every 12^th^ section was counted. There was no correction factor applied to the values reported. Only one ovary per animal was used.

Cultured ovaries, used for histological sections followed by immunostaining, were fixed in 4% paraformaldehyde/PBS over night at 4°C. After 70% ethanol washes, ovaries were embedded in paraffin and serially sectioned at 5μm. These ovaries were immunostained using standard methods. Briefly, slides were deparaffinized and re- hydrated prior to antigen retrieval using sodium citrate buffer. Slides were blocked with 5% goat serum (PBS/Tween 20) and incubated at 4°C overnight with primary antibodies: mouse anti-p63 (1:500, 4A4, Novus Biologicals); and rabbit anti-MVH (1:1000, Abcam). Afterwards, sections were incubated with Alexa Fluor(r) secondary antibodies for one hour and Hoechst dye for 5 minutes. Slides were mounted with ProLong Anti-fade (Thermo-Fisher) and imaged.

Histological images present in the manuscript were obtained from slides digitized using a Leica Scanscope CS2.

### Immunofluorescence of meiotic chromosome surface spreads

Meiotic surface spreads of prophase I female meiocytes were prepared using a drying-down technique ^9^. Meiotic stages were determined based on SYCP3 staining patterns. Slides were stored at -80?C until stained. For staining, slides were brought to room temperature (RT) and washed once with PBS+0.1% Tween-20 (PBS-T). Slides were blocked for 40 minutes at RT with PBS-T containing 5% normal goat serum (5%GS-PBS-T). Primary antibodies were diluted into 5%GS-PBS-T and incubated overnight at RT in a humidified chamber. Antibodies and dilutions used included: rabbit anti-RAD51 (1:250 abcam ab 176458), mouse anti-SYCP3 (1:600 Abcam) and guinea pig anti-HORMAD2 antibody (1:1000, kind gift from Attila Toth). Secondary antibodies used were diluted 1:1000 in in 5%GS-PBS-T and included goat anti-rabbit Alexa 488/594, goat anti-mouse Alexa 488/594 and goat anti-guinea pig Alexa 488/594.

Images were taken using an Olympus microscope with 40X lens or 100X immersion oil lens and CCD camera.

### Focus Quantification

Foci were quantified both manually, through the visualization and annotation of individual foci, and also semi-automatically using Fiji-ImageJ ^10^. Semi-automated counts were performed using binary images obtained from the RAD51-labeled channel, with the threshold set above background level. The count was obtained after performing “Watershed”, by the “Analyze Particles” functionality with size set for 1.5 to infinity. Cell counts that displayed discrepancy of more than 20% between manual and semi-automated counts were discarded.

### Statistical analysis

All statistical analyses were done using JMP Pro12 software (SAS Inc., Cary, NC-USA, version 12.0.1). Comparisons of fertility and follicle counts between genotypic groups were tested using both the Tukey honest significance different (HSD) and the non-parametric, one-way ANOVA test (Kruskal -Wallis). Both tests provided concordant results. RAD51 focus counts were analyzed using a mixed model with animal ID as random effect and genotype as fixed effect. Least square means (LSMeans) differences were tested using Tukey HSD.

### Fertility Test

To test if HORMAD2 deficiency was able to rescue the *Trip13*^*Gt/Gt*^sterility phenotype, three double mutant females were mated to wild type C3H/HeJ males proven to be fertile through previous matings. Each female provided more than 4 consecutive litters up to the time of preparation of this manuscript. All three females originated from different litters. *Trip13*^*Gt/Gt*^littermates were housed with fertile males and used as negative controls.

---

**Supplementary Fig. 1.**
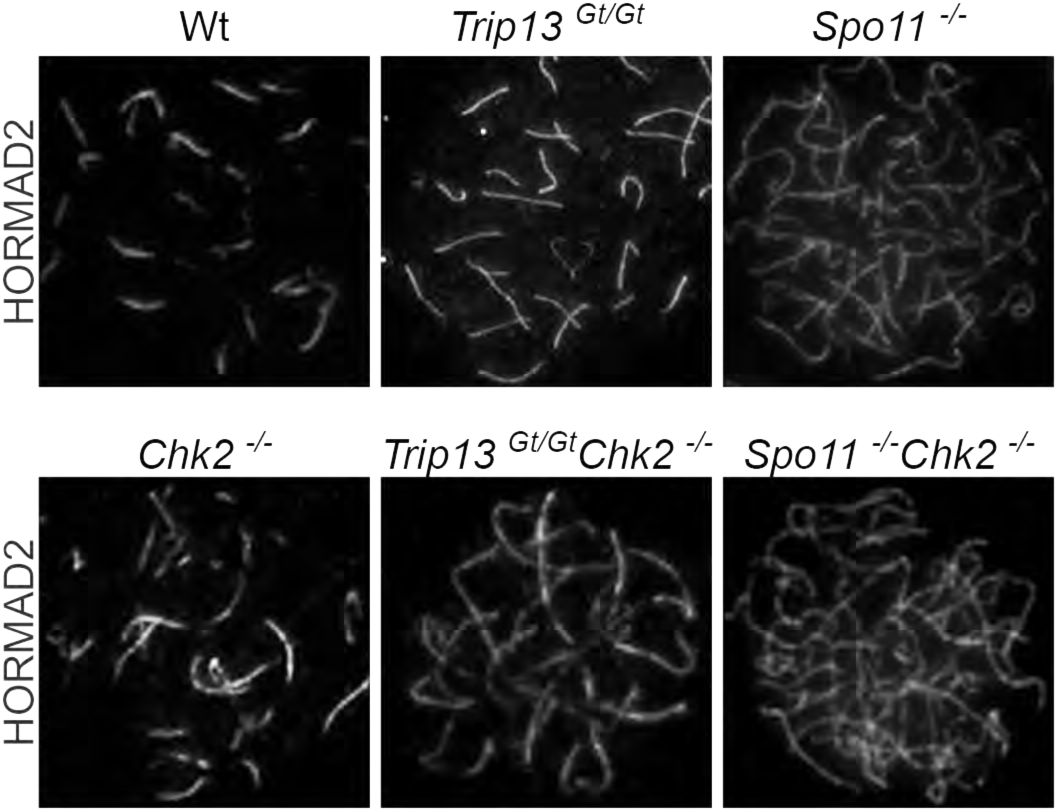
CHK2 is not required for HORMAD localization to meiotic chromosome axes of Spo11-*1*-or Trip13Gt!Gt oocytes. Shown are zygo tene and zygotene-like nuclei from indicated geno types.

**Supplementary Fig. 2.**
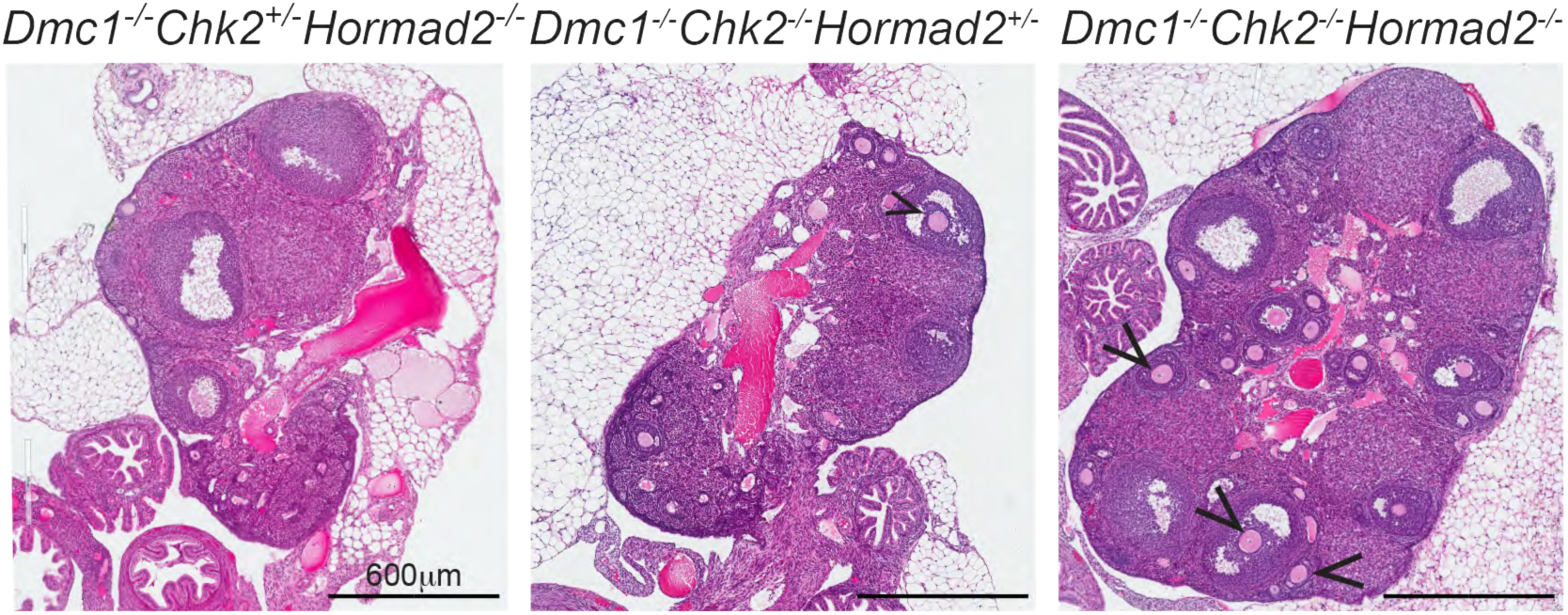
H&E stained histological sections of 8 week old ovaries of specified genotypes. Black arrowheads indicate antral follicles.

**Supplementary Fig. 3.**
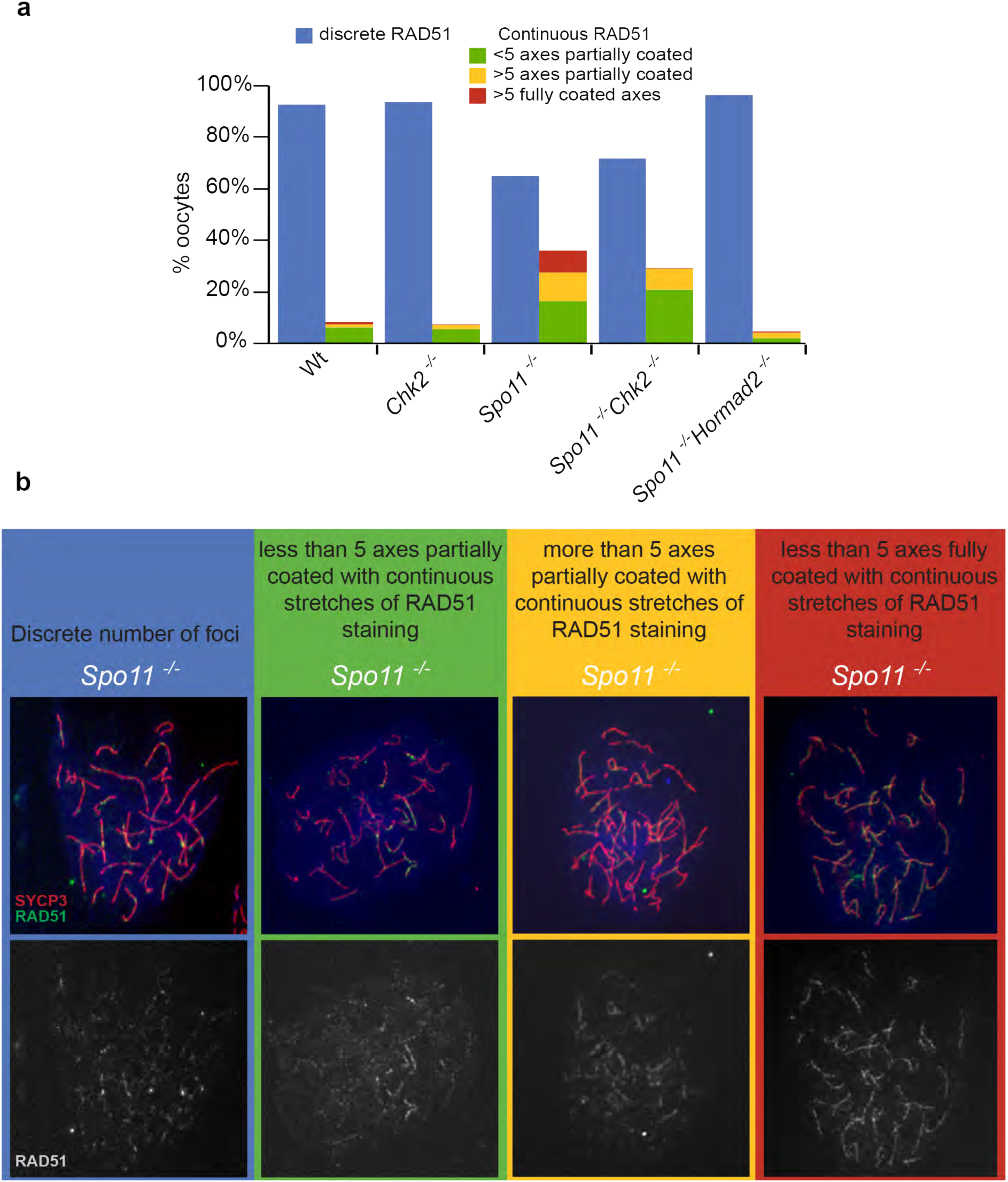
Patterns of RAD51 staining on oocyte meiotic chromosomes of various genotypes. (a) The plots shows the percentage of oocytes from the specified genotypes, color coded for either discrete foci or varying levels of continuous staining patterns. (b) Classification of the different levels of continuous RAD51 immunostaining. All chromosome spreads are derived from newborn mice. RAD51 quantification was performed in images derived from an objective with 0.45μm resolving power. At this resolution, the RAD51 signals could be classified as discrete or continuous (coating AEs and/or SCs). It is likely that these continuous staining regions consist of numerous distinct foci. Because these were not enumerated in the calculation of discrete SPO11-independent DSBs, the actual number of SPO11-independent DSBs is probably higher than reported here.

**Supplementary Fig. 4.**
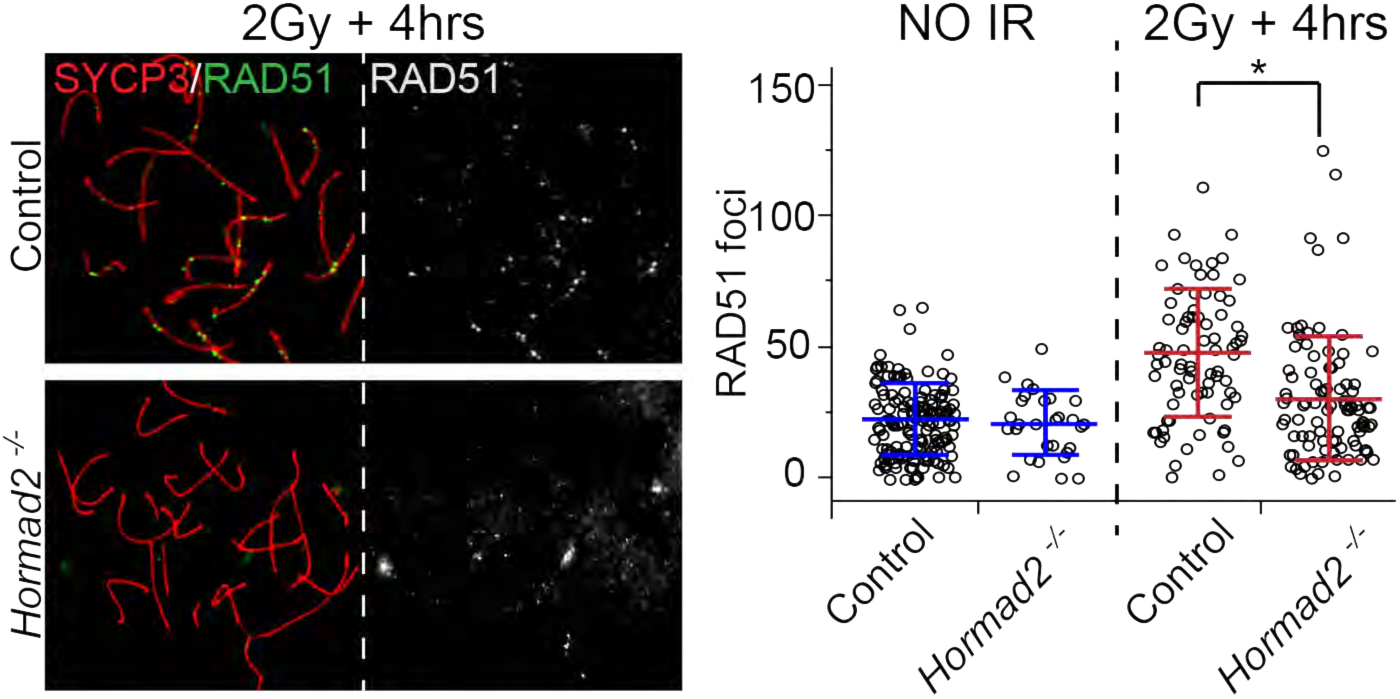
Hormad2^-1^ oocytes exhibit accelerated repair of radiation-induced DSBs. IR= ionizing radiation.

